# Fate mapping caudal lateral epiblast reveals continuous contribution to neural and mesodermal lineages and the origin of secondary neural tube

**DOI:** 10.1101/045872

**Authors:** Aida Rodrigo Albors, Pamela A. Halley, Kate G. Storey

**Author notes:** equal contribution (Aida Rodrigo Albors, Pamela A. Halley). Correspondance: Kate G. Storey School of Life Sciences University of Dundee Dow Street Dundee, DD1 5EH.

## Abstract

The ability to monitor and manipulate epiblast cells in and around the primitive streak of the mouse embryo is important for investigating how cells maintain potency and how distinct cell fates are established. Here, we report development of a key resource for such studies, a mouse line in which a tamoxifen-inducible Cre recombinase construct replaces an endogenous gene, *Nkx1.2*, whose expression demarcates the epiblast adjacent to and including the primitive streak. We show that this Nkx1.2CreER^T2^ line drives transgene expression in the endogenous *Nkx1.2* domain. Labelling this caudal epiblast cell population at embryonic day (E) 7.5 confirmed contribution to all three germ layers at later stages. Labelled cells were also retained within the caudal lateral epiblast at E8.5 and E9.5 and E10.5 tailbud. A subset of these cells co-expressed early neural (Sox2) and mesodermal (Bra) markers. These findings support the existence of neuromesodermal progenitors within the *Nkx1.2* cell lineage. Consistent with the retention of such bipotent progenitors throughout body axis elongation, labelling *Nkx1.2-*expressing cells at E10.5 and assessment at E11.5 demonstrated continued contribution to both neural and mesodermal lineages. Furthermore, detailed analysis of *Nkx1.2* expression revealed a novel domain in tailbud mesenchyme that is contiguous with neural tissue. The presence of labelled Sox2/Bra co-expressing cells in this mesenchyme cell population suggests that this is the retained neuromesodermal progenitor pool, which gives rise to both new paraxial mesoderm and new neural tissue, generated now via secondary neurulation. Here, we introduce this caudal-most *Nkx1.2* domain as the neuromesodermal lip.

## Introduction

Epiblast cells in early mouse embryos contribute to all three germ layers, endoderm, mesoderm, ectoderm/neuro-ectoderm and this is gradually restricted as development proceeds. Once the primitive streak has formed, adjacent epiblast cells, known as the caudal lateral epiblast (CLE), appear to contribute largely to neural and mesodermal lineages. Evidence for this comes from retrospective clonal analysis, which predicts that some CLE cells become neuro-mesodermal progenitors (NMps) that then persist during body axis elongation (Henrique et al., 2015; Tzouanacou et al., 2009) and also from homotopic grafting of groups of epiblast cells (Wilson et al., 2009; Wymeersch et al., 2016). Some NMps are also located more medially in the node streak border (NSB), which additionally includes presumptive notochord cells (Cambray and Wilson, 2002, 2007; Tsakiridis et al., 2014; Wymeersch et al., 2016). Unique markers for NMps have yet to be identified, but co-expression of early neural and mesodermal markers, Sox2 and Brachyury (Bra) correlates well with the predicted location of these cells in the embryo (Henrique et al., 2015; Tsakiridis et al., 2014; Wymeersch et al., 2016). Furthermore, embryonic stem (ES) cell derived cells co-expressing these genes readily differentiate into neural or mesodermal lineages in vitro (Gouti et al., 2014; Turner et al., 2014) and this appears to be the case at the single cell level (Tsakiridis and Wilson, 2015). Neural tissue generated by epiblast close to the primitive streak contributes to posterior hindbrain and spinal cord (Forlani et al., 2003; Lawson et al., 1991; Lawson and Pedersen, 1992). However, about mid-way through body axis elongation amniote embryos form a tailbud and subsequent neural tissue is now produced by secondary neurulation of a core of condensed mesenchyme (Schoenwolf, 1984). Tailbud formation involves internalisation of remnants of the primitive streak, however, the precise fate of the CLE and the role(s) of its derivatives in the continued generation of neural and mesodermal tissue is poorly understood.

The ability to manipulate gene expression in the CLE will allow detailed fate mapping of this cell population as well as facilitate investigation of the bipotent cell state and the mechanisms that establish neural and mesodermal differentiation programmes in vivo. The latter involves the coordinated onset of tissue specific differentiation genes and a change in cell behaviour as cells undergo epithelial to mesenchymal transition to become mesoderm. Here we report development of a key resource for such studies, a mouse line in which a tamoxifen-inducible Cre recombinase construct replaces an endogenous gene whose expression demarcates the CLE.

The NK-1 homeobox 2 transcription factor, *Nkx1.2*, also known as *Sax1*, is the earliest known CLE expressed gene and it distinguishes this epiblast cell population from differentiating neural and mesoderm tissue (Spann et al., 1994). *Nkx1.2* is a homologue of the *Drosophila NK-1/S59* gene, which is expressed in ganglion mother cells, transit-amplifying cells of the fly central nervous system that generate neurons or glia, and also in muscle founder cells and their progenitors as well as midgut cells (Bate, 1990; Dohrmann et al., 1990). *Nkx1.2* is widely conserved across species; representatives are present in chordates and helminthes as well as many vertebrates. In the mouse, a paralogous gene *Nkx1.1 (Sax2*) has been identified (Bober et al., 1994; Simon and Lufkin, 2003) and two closely related Nkx1 genes have been described in the Zebrafish (Bae et al., 2004).

The expression pattern of *Nkx1.2* has been characterised in chick, mouse and Zebrafish (Bae et al., 2004; Rangini et al., 1989; Schubert et al., 1995; Spann et al., 1994). In the chick, *Nkx1.2* (originally named *cHox3* and later *cSax1*) is detectable at early primitive streak stages ((Hamburger and Hamilton, 1951) HH 2), but is only visualised by mRNA in situ hybridisation after formation of the first few somites, when it appears in the CLE/stem zone (Delfino-Machin et al., 2005; Rangini et al., 1989; Spann et al., 1994). This earliest expression then persists in the CLE and expands slightly rostral into the preneural tube (adjacent to presomitic mesoderm) during body axis elongation (Delfino-Machin et al., 2005). Later phases of expression are not characterised in detail in the chick, but include a subset of interneurons in the spinal cord/hindbrain and expression in the ventral midbrain by HH15 (Schubert and Lumsden, 2005).

In the mouse, *Nkx1.2* is first detected at E7 in the forming CLE next to the primitive streak, is retained until almost the end of axial elongation (E12.5) and is consistently down-regulated in neural tissue adjacent to the most recently formed somite (Schubert et al., 1995). A later phase of *Nkx1.2* expression is detected from stages (E10-10.5) in bilateral ventral stripes of interneurons in more rostral spinal cord as well as in rhombomere 1. At this time, *Nkx1.2-* expressing cells also appear in discrete cell populations in the developing brain, including the ventral midbrain (Schubert et al., 1995) where, in the chick, *Nkx1.2* specifies the cells contributing to the medial longitudinal fascicle (Schubert and Lumsden, 2005).

Regulation of *Nkx1.2* in the CLE has been studied in chick and mouse embryos. In the chick, removal of notochord, somites or presomitic mesoderm does not alter *Nkx1.2* expression assessed at a late time point when new presomitic mesoderm, but not notochord or somites has been regenerated (Spann et al., 1994). Subsequent work showed that signals from the presomitic mesoderm promote *Nkx1.2* onset as the CLE/stem zone forms and that the FGFR/Erk pathway is required for maintenance of *Nkx1.2* in CLE cells; however, FGF alone is insufficient to induce *Nkx1.2* in this cell population (Delfino-Machin et al., 2005; Diez del Corral et al., 2002). Consistent with these data, *Nkx1.2* expression is lost in mice mutant for the early mesodermal gene *Bra*, which lacks caudal presomitic mesoderm (Schubert et al., 1995). The discrete downregulation of *Nkx1.2* as neural differentiation commences is regulated by signals from the more rostral differentiating presomitic mesoderm (Diez del Corral et al., 2002). This likely reflects the action of retinoic acid, which is synthesized in this tissue and inhibits *Fgf8* expression in the stem zone/CLE (Diez del Corral et al., 2003; Kumar and Duester, 2014; Sirbu and Duester, 2006). Intriguingly, recent work shows that *Nkx1.2* in turn promotes *Fgf8* transcription (Sasai et al., 2014), further linking this transcription factor to signalling pathways that regulate differentiation along the body axis. Consistent with this, inhibition of *Nkx1.2* leads to precocious expression of neural differentiation genes including *Pax6* (Sasai et al., 2014). However, *Pax6* mutants exhibit normal *Nkx1.2* expression indicating that reciprocal repression by Pax6 is not required to restrict *Nkx1.2* to the CLE (Schubert et al., 1995). As noted above, many cells in the CLE/Nkx1.2 domain co-express Sox2 and Bra (Delfino-Machin et al., 2005; Olivera-Martinez et al., 2012; Tsakiridis and Wilson, 2015) and this bipotent cell state can be induced in differentiating mouse and human ES cells by a combination of FGF and Wnt signalling (Gouti et al., 2014; Tsakiridis et al., 2014) (reviewed in (Henrique et al., 2015)). Such CLE-like cells also express *Nkx1.2* (Gouti et al., 2014; Tsakiridis et al., 2014) and so it is likely that in the embryo Wnt signalling together with FGF promotes *Nkx1.2*. This transcription factor has further been shown to induce Bra expression in P19 cells in part by repressing expression of *Tcf3* (Tamashiro et al., 2012); a step that augments Wnt induction of *Bra* (Marikawa et al., 2009; Tanaka et al., 2011) and so further links *Nkx1.2* to maintenance of this bipotent state.

Here, we present a detailed description of endogenous *Nkx1.2* expression in the mouse embryo and the creation and characterisation of a mouse line in which tamoxifen-inducible Cre recombinase is cloned in place of the endogenous *Nkx1.2* gene. We demonstrate that this line can be used to manipulate gene expression in the CLE/tailbud and provide a detailed fate map of these cell populations. This includes a novel *Nkx1.2* domain in the tailbud mesenchyme, which contains cells co-expressing neural and mesodermal genes and that contributes to both neural and mesodermal lineages.

## Results and Discussion

### *Nkx1.2* is expressed in the caudal epiblast from E7 and within the tailbud from E10.5 to E11.5

To document in detail *Nkx1.2* expression in developing mouse embryos, we carried out RNA in situ hybridisation in whole embryos throughout the period of body axis elongation. We then localised transcripts to specific cell populations in transverse sections. In agreement with a previous report (Schubert et al., 1995), *Nkx1.2* transcripts were first detected at E7-7.5 in the epiblast in the region of and adjacent to the node and the primitive streak, known as the CLE; including posterior primitive streak (Figure 1A-C). At E8.5 *Nkx1.2* expression persisted in these cell populations, but was now also detected in epiblast cells in the open neural plate anterior to the node (Figure 1D, E). *Nkx1.2* transcripts were present in cells at the ventral midline in the region of the node (Figure 1Eb), but rapidly downregulated and lacking in the floor plate of the neural plate (Figure 1Ea). By E9.5 the most anterior *Nkx1.2*-expressing cells have begun to form a neural tube, with expression strongest dorsally (Figure 1F, G). In the caudal-most part of this domain, transcripts remain confined to the epiblast cell layer and are not detected within the primitive streak (Figure 1Gb, Gc). At E10.5 *Nkx1.2* persists caudally in the closed neural tube (Figure 1H, I), but transcripts are now also detected in the tailbud mesenchyme that is contiguous with the caudal-most neural tube (Figure 1Ic). Intriguingly, the appearance of this novel *Nkx1.2* domain coincides with the switch to formation of neural tissue by secondary neurulation (Schoenwolf, 1984). At E11.5 *Nkx1.2* transcripts continue to be detected in the caudal-most neural tissue and contiguous mesenchyme of the tailbud (Figure 1J, K). By E12.5, when elongation of the tail is coming to a halt, *Nkx1.2* expression is lost (Figure 1L, M). *Nkx1.2* is also detected outside the caudal region in a sub-population of motor neurons in the hindbrain and spinal cord at E10.5 (Figure 1N) and in the medial longitudinal fascicle of the midbrain (Schubert et al., 1995) (Figure 1O, Oa). Taken together, these data show that *Nkx1.2* is a consistent marker of the CLE and newly generated caudal neural progenitors throughout body axis elongation. An intriguing possibility is that the novel *Nkx1.2* domain found in tailbud mesenchyme may additionally identify the cellular source of the secondary neural tube.

**Figure 1.**
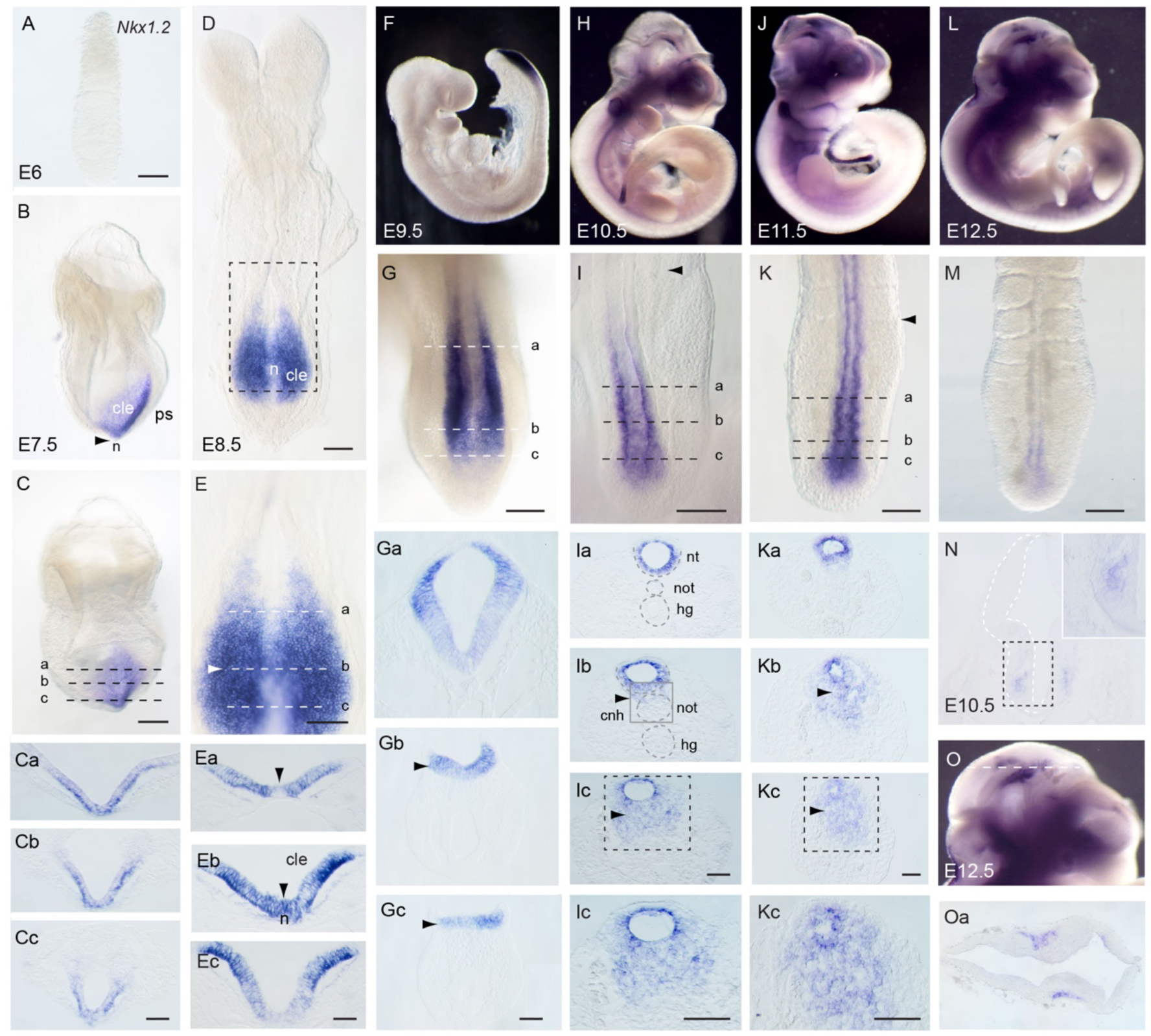
Expression of *Nkx1.2* in the developing mouse embryo. (A) *Nkx1.2* transcripts were not detected at E6.0. (B) Lateral and (C) posterior views of an E7.5 embryo (early head fold), where *Nkx1.2* transcripts were detected in the CLE (cle) adjacent to the node (n) and primitive streak (ps). (Ca-Cc) Transverse sections (TS) through the regions indicated in C. (D) E8.5 embryo (8-10 somites) and (E) higher magnification of the caudal end of the embryo. (Ea-c) TS through the regions indicated in E. (Ea) *Nkx1.2* transcripts were detected in the epiblast except in the ventral midline rostral to the node. (Eb-c) More caudally, *Nkx1.2* is expressed across the epiblast and lost in the deep layers of primitive streak. (F) E9.5 embryo and (G) dorsal view of end of the embryo. (Ga-c) TS through the regions indicated in G. (Ga) *Nkx1.2* transcripts were only detected in the closing neural tube and (Gb-Gc) caudal epiblast but not within the primitive streak. (H) E10.5 embryo and (I) dorsal view of the tail tip. *Nkx1.2* expression is lost at the level of most recently formed somite (black arrowhead). (Ia-c) TS through the regions indicated in I. (Ia) *Nkx1.2* was only expressed in the closed neural tube (nt) at more rostral levels. (Ib) In the CNH (cnh) region (where the caudal end of the notochord meets the neural tube), *Nkx1.2* transcripts were detected in neural tube, but also in associated mesenchymal cells. (Ic) Caudal to the CNH, *Nkx1.2* transcripts were abundant in tailbud mesenchyme. (J) E11.5 embryo and (K) dorsal view of the tail tip. *Nkx1.2* was expressed in the most recently formed secondary neural tube and contiguous tailbud mesenchyme. (Ka-c) TS through the regions indicated in K. (Ka) *Nkx1.2* was only expressed in the neural tube rostral to the CNH (Kb-c). Caudal to the CNH, and similar to E10.5, *Nkx1.2* was transcribed in the forming neural tube but also in a broad tailbud mesenchyme domain. (L) E12.5 embryo and (M) dorsal view of the tail tip. At this stage, *Nkx1.2* signal starts to decrease in the tailbud. (N) *Nkx1.2* expression in a subpopulation of motor neurons of the hindbrain/spinal cord of an E10.5 embryo. (O) Higher magnification of L showing *Nkx1.2* in the medial longitudinal fascicle of the midbrain. (Oa) TS through the region indicated in O. Scale bars in wholemounts, 100 μm; scale bars in TS, 50 μm

### Generation of Nkx1.2CreER^T2^ mouse line

Prompted by these observations, we sought to investigate the contribution of *Nkx1.2-* expressing cells to the developing mouse embryo. To do so, we generated a mouse in which we could manipulate gene expression in *Nkx1.2*-expressing cells in a temporally-controlled manner, by driving CreER^T2^ recombinase under the control of the endogenous *Nkx1.2* locus. The CreER^T2^ sequence was knocked in to this locus using standard targeting methods in C57BL/6 ES cells by Taconic Biosciences. The DNA constructs used are summarised in Figure 2. Two breeding pairs of C57BL/6-Nkx1.2^tm2296(Cre-ER(T2))Arte^ (Nkx1.2CreER^T2^) heterozygous mice were provided by Taconic. These mice carried a puromycin-expressing cassette flanked by FLP sites, which was removed upon crossing to Flp-expressing mice. The resulting animals were then bred to the homozygous condition to establish a breeding colony.

**Figure 2.**
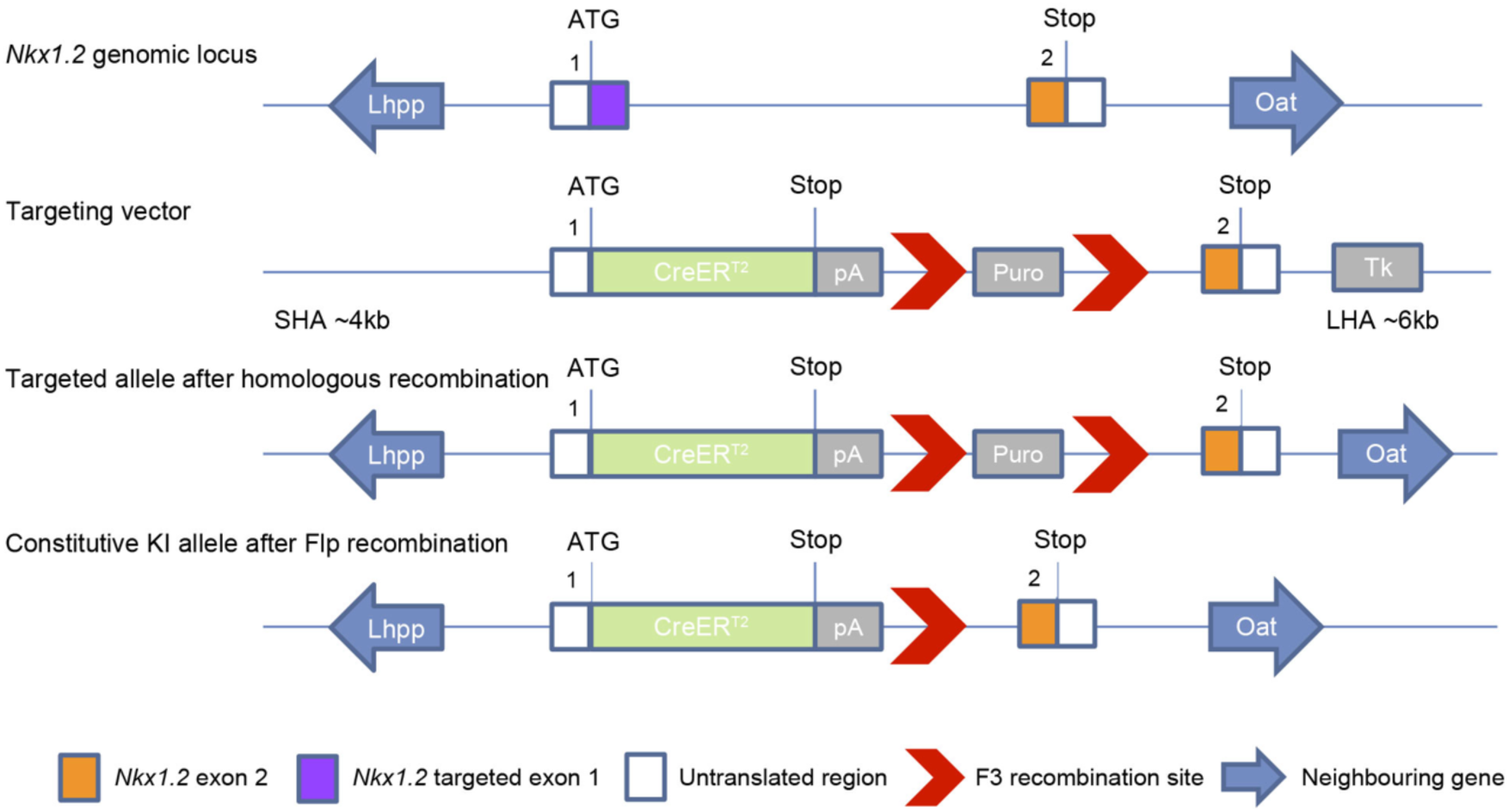
Strategy to knock-in the CreER^T2^ cassette into the *Nkx1.2* genomic locus. Targeting vector designed to insert into the downstream region of exon 1 of the mouse Nkx1.2 locus by homologous recombination. Positive clones were injected into mouse blastocysts and transferred to mice. The resulting chimeric mice were bred to Flp Deleter mice that ubiquitously express Flp recombinase to remove the puromycin selection marker to give rise to the Nkx1.2CreER^T2^ mouse line. SHA, short homologous arm; LHA, long homologous arm; KI, knock-in.

Next, to label *Nkx1.2*-expressing cells in the developing embryo, homozygous Nkx1.2CreER^T2^ females were mated with heterozygous or homozygous males harbouring a loxP-flanked STOP sequence upstream of a YFP reporter gene under the control of the ubiquitous *ROSA26* promoter (ROSA26-floxed-stop-YFP knock-in mice) (Srinivas et al., 2001). In the resulting Nkx1.2CreER^T2^ floxed YFP mice (Nkx1.2CreER^T2^/YFP), upon tamoxifen administration, Cre-mediated recombination will result in the deletion of the *loxP*-flanked STOP sequence and expression of the YFP reporter in *Nkx1.2*-expressing cells.

*Nkx1.2* has been knocked out in the mouse, but did not generate a phenotype in either heterozygous or homozygous animals (Frank Schubert and Peter Gruss pers. comm.). This is most likely due to functional redundancy with the related, paralogous *Nkx1.1* gene (Bober et al., 1994). We substantiate this unpublished finding here, with the maintenance of a homozygous Nkx1.2CreER^T2^ mouse line for at least 8 generations presenting no deleterious phenotype.

### Nkx1.2CreER^T2^ drives transgene expression in the endogenous *Nkx1.2* domain

To assess how faithfully the Nkx1.2CreER^T2^ transgene recapitulates endogenous *Nkx1.2* expression, timed-pregnant Nkx1.2CreER^T2^/YFP mice were exposed to tamoxifen by oral gavage at E6.75 and embryos assessed for YFP expression at E7.5. This showed YFP-labelled epiblast cells in the region of the node and primitive streak (Figure 3A, B) and so was consistent with the pattern of endogenous *Nkx1.2* transcripts at ~E7.5 (Schubert et al., 1995) (Figure 1B, C). To assess potential spontaneous CreER^T2^ recombination in Nkx1.2CreER^T2^ mice, we next looked at YFP expression in the absence of tamoxifen induction in E8.5 embryos. All these embryos showed low levels of spontaneous recombination, from 4 to 9 YFP+ cells per embryo (6 ± 1 cells/embryo, mean and s.e.m.) (Figure 3C). The close proximity and number of YFP+ cells suggests that they originate from one recombination event; given that it takes ~ 4h to detect YFP, that cells have a cell cycle of ~6-7h (Snow, 1977; Tzouanacou et al., 2009), this would be 2 or 3 divisions following labelling of a single cell at ~E7.5. Importantly, the labelled cell group was always within canonical *Nkx1.2*-expressing regions. It is thus possible that cytoplasmic CreER^T2^ does on rare occasions translocate into the nucleus in the absence of tamoxifen. However, because only *Nkx1.2*-expressing cells produce CreER^T2^, this infrequent spontaneous recombination event within the embryo may in fact be useful for clonal lineage tracing of these cells. Overall, these data show that upon tamoxifen administration, CreER^T2^-mediated recombination at the *Nkx1.2* locus leads to faithful YFP-labelling of *Nkx1.2*-expressing cells.

**Figure 3.**
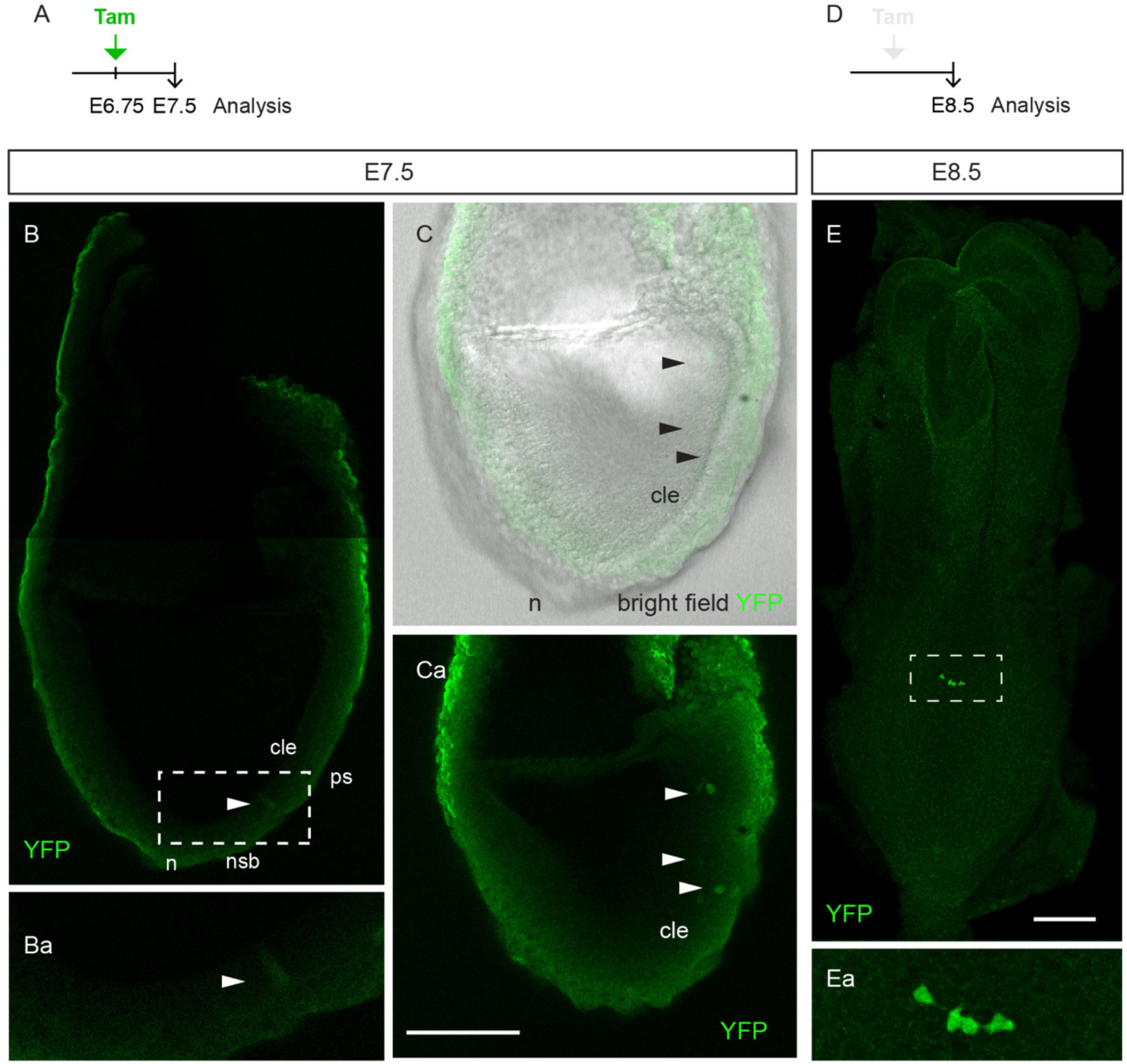
Tamoxifen induction drives CreER^T2^-mediated recombination specifically in *Nkx1.2*-expressing cells. (A) Timed-pregnant Nkx1.2CreER^T2^/YFP mice received tamoxifen at E6.75 and embryos analysed for YFP-labelled cells at E7.5 in wholemount. (B) Parasagittal single optical section through a late bud stage Nkx1.2CreER^T2^/YFP embryo (~E7.5). YFP-labelled cells were present in the node-streak border (nsb) region. (Ba) Higher magnification of the region indicated in B. (C) Maximum intensity projection (MIP) of 4 optical sections of the embryo in B. YFP+ cells were also evident throughout the CLE (arrowheads). (Ca) YFP channel of B. (Low numbers of recombined cells following exposure at E6.75 may reflect inaccessibility of the embryo to tamoxifen at this early stage as well as onset of Nkx1.2 transcription only from at E7.0)(D) Nkx1.2CreER^T2^/YFP embryos that were not exposed to tamoxifen were analysed at E8.5. (E) Confocal MIP of an Nkx1.2CreER^T2^/YFP E8.5 embryo in the absence of tamoxifen induction. A small group of YFP+ cells was found in the CLE. (Ea) Higher magnification of the region indicated in C. Labels are as in Figure 1. Scale bars, 100 μm.

### Early *Nkx1.2*-expressing epiblast contributes to specific cell populations in all three germ layers

To follow *Nkx1.2*-expressing cells and their progeny throughout body axis elongation, pregnant Nkx1.2 CreER^T2^-YFP mice were exposed to a single dose of tamoxifen at E7.5 and the contribution of YFP^+^ cells was assessed in embryos at different developmental stages (Figure 4A). Assessment of embryos at E8.5 (n=7 embryos) revealed scattered single cells across the presumptive midbrain/anterior hindbrain as the rostral-most limit of YFP^+^/CLE-derived cells (Figure 4B, Ba). YFP+ cells were consistently absent from the first 4-5 somites and from the notochord (Figure 4B, Bc), but were then found contiguously in the neural tube beginning in the presumptive posterior hindbrain (Figure 4B, Bb) and then, more caudally, throughout the CLE and primitive streak (Figure 4B, Bd-Bg). YFP+ cells were also present in the recently ingressed mesoderm at the node-streak border and in the presomitic mesoderm emerging from the primitive streak (Figure 4Bd-Bg).

**Figure 4.**
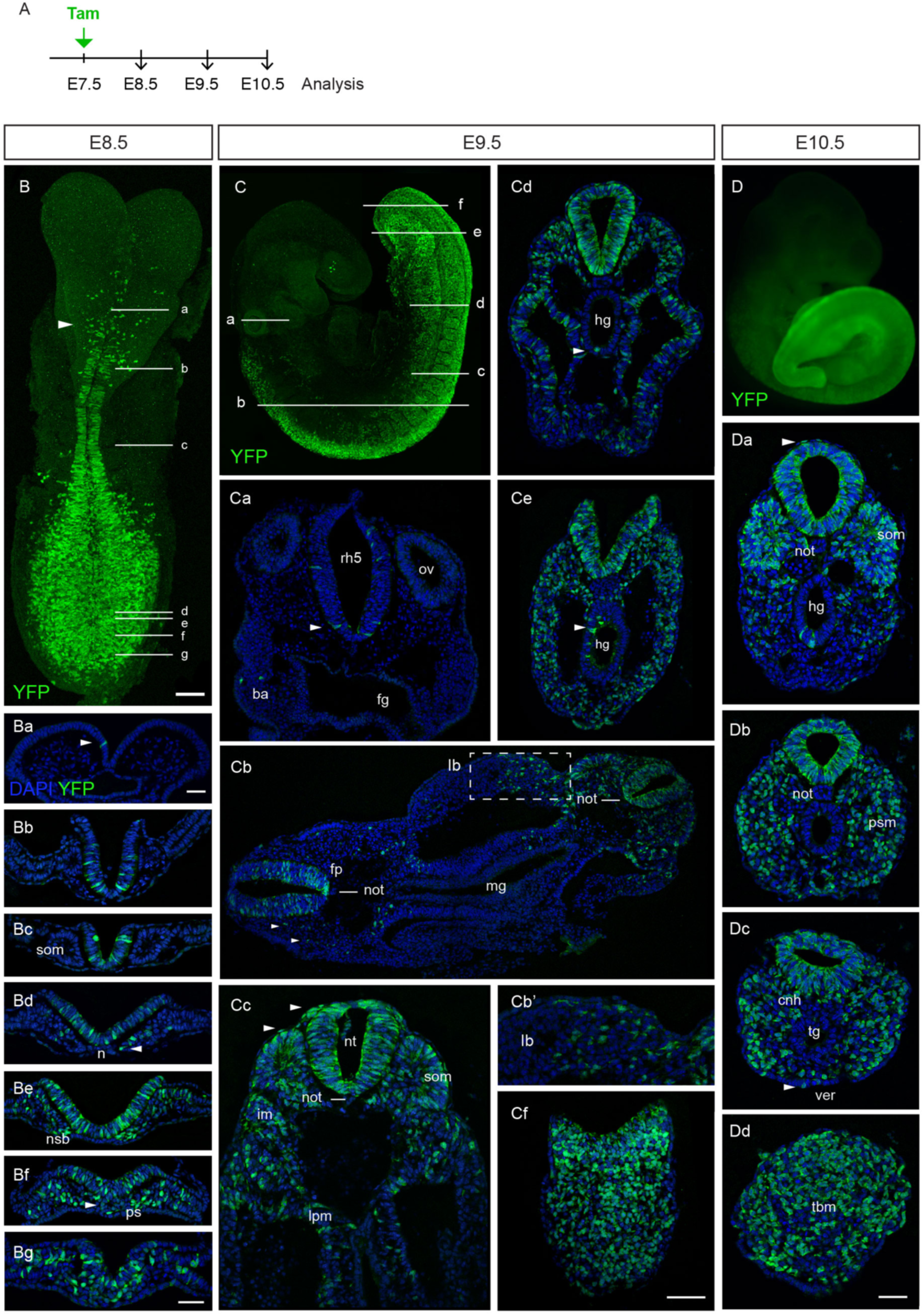
Descendants of E7.5 *Nkx1.2*-expressing cells in the developing mouse embryo. (A) Timed-pregnant Nkx1.2CreER^T2^ mice received tamoxifen at E7.5 and the contribution of YFP+ cells to developing embryos assessed at E8.5, E9.5, and E10.5. (B) Confocal maximum intensity projection (MIP) of an E8.5 embryo labelled with an antibody against YFP. The arrowhead marks the presumptive midbrain/anterior hindbrain boundary. (Ba-Bg) Transverse Sections (TS) through the regions indicated in A. (C) Confocal MIP of a E9.5 embryo stained against YFP. (Ca) TS through the otic vesicles (ov) and rhombomere 5 (rh5). Scattered cells were found mainly in ventral neural tube at this level (arrowhead). A few labelled cells populated the branchial arches (ba). The foregut (fg) was, however, always unlabelled. (Cb) TS comprising the rostral (left) and caudal (right) levels of the trunk, as indicated in B. YFP^+^ cells located mostly in the floor plate (fp) of the neural tube at more rostral levels, but spanned the whole dorsoventral axis of more caudal neural tube. YFP+ cells generated neural crest cells (arrowheads), contributed to somites caudally, and to limb bud (lb) mesenchyme. YFP+ cells were absent from notochord (not) and midgut (mg). (Cb’) Higher magnification of limb bud mesenchyme in Bb (dashed box). (Cc) TS approximately through the region indicated in C. YFP+ cells contributed to the neural tube (nt), somites (som), intermediate mesoderm (im), lateral plate mesoderm (lpm), and surface ectoderm (arrowheads). (Cd, e) TS through the regions indicated in C. YFP+ cells were found frequently in the hindgut (hg) (arrowhead). (Cf) TS through the region indicated in C. YFP+ cells extended to the caudal neural plate and underlying mesenchyme. (D) Widefield fluorescence image of an E10.5 embryo that received tamoxifen at E7.5. Most of the caudal half of the body derived from YFP+ cells at this stage. (Da) TS at tail somite levels. YFP+ cells made most of the neural tube and somites (som) and also contributed to hindgut (hg) endoderm and surface ectoderm (arrowhead), but were absent from the notochord (not). (Db) TS caudal to Da showed labelled unsegmented presomitic mesoderm (psm). (Dc) TS caudal to Db, across the CNH (cnh). Besides neural tube and paraxial mesoderm, YFP+ cells occupied a small region between the ventral neural tube and condensing notochord. YFP+ cells were also present in the VER (ver). (Dd) TS through the tailbud mesenchyme (tbm). The images are representative images of each stage and rostrocaudal level. A summary of tissue contributions and number of embryos analysed at each developmental stage can be found in Table 1. Scale bars, 50 μm.

A day later, in E9.5 embryos, again a few scattered cells were located in the midbrain and also the roof of the anterior hindbrain and as well as the developing eye (Figure 4C). The rostral-most limit of contiguous YFP labelling was now clearly located in the hindbrain just rostral to the otic vesicle in rhombomere 5 (Figure 4C). More caudally, YFP+ cells were concentrated ventrally, including in the floor plate of the neural tube (Figure 4Ca, Cc). At forelimb levels, YFP+ cells were found throughout the dorsoventral extent of the neural tube as well as in paraxial mesoderm (somites) and its derivatives, including mesenchyme cells migrating in to the limb bud (Figure 4Cb, Cc). YFP+ cells also contributed extensively to intermediate and lateral plate mesoderm (Figure Cc). From forelimb levels, labelled cells also appeared in the surface ectoderm (Figure 4Cc-f), and streams of YFP+ neural crest cells emanated from the dorsal neural tube (Figure 4Cc). In line with embryos at E8.5, YFP+ cells were also absent from the first 4-5 somites of E9.5 embryos, but contributed to both medial and lateral compartments of the caudal-most 11-12 somites (Figure 4C, Cb, Cd). Labelled cells were absent from the notochord (Figure 4Ca-c), although a few isolated YFP+ cells were found in this tissue in one embryo. In all E9.5 embryos examined, YFP+ cells were absent from the fore-and midgut (Figure 4Ca-c), but frequently found in the hindgut (Figure 4Cd, Ce). At the caudal end of the embryo, YFP^+^ cells made most of the caudal neuropore and underlying mesenchyme, and also again contributed to the surface ectoderm (Figure 4Cc-f).

Taken together, these findings suggest that the majority of descendants of E7.5 *Nkx1.2-* expressing cells generate neural tube and trunk mesoderm (except axial mesoderm), and that some cells labelled at this early stage contribute to surface ectoderm and to the hindgut endoderm (Table 1). The latter is consistent with direct fate mapping of single cells (Lawson et al., 1991; Lawson and Pedersen, 1992) and of cell groups (Tam and Beddington, 1987; Wilson and Beddington, 1996) in the early primitive streak which all indicate that the gut endoderm lineage is derived from primitive streak epiblast prior to E8.5.

**Table 1.** Tissue contribution of *Nkx1.2*-expressing cells labelled at E7.5 to E9.5 and E10.5 embryos. The numbers for each tissue type show the percentage of embryos in which labelled cells where found in particular tissues, assessed in transverse sections.

| embryonic stage | no. of embryos | ectoderm |  | endoderm |  |  | mesoderm |  |  |  |
| --- | --- | --- | --- | --- | --- | --- | --- | --- | --- | --- |
|  |  | neuroectoderm | surface ectoderm | foregut | midgut | hindgut | axial mesoderm (not) | paraxial mesoderm | intermediate mesoderm | lateral plate mesoderm |
| E9.5 | 4 | 100 | 100 | 0 | 0 | 100 | 25 | 100 | 100 | 100 |
| E10.5 | 7 | 100 | 100 | 0 | 0 | 100 | 14 | 100 | 100 | 100 |

Between E9.5-10.5, axial progenitors complete trunk formation and begin tail formation. As contribution patterns at E9.5 are largely elaborated in the body axis at E10.5, we focused here on the E10.5 tailbud region. Here most of the caudal body derives from cells that expressed *Nkx1.2* at E7.5, including cells in the neural tube and paraxial mesoderm, with occasional cells in the hindgut and surface ectoderm, and very rarely in axial mesoderm (notochord) (Figure 4D, Da-c) (Table 1). Formation of the tailbud involves the ventral bending of the tail fold, which brings the posterior primitive streak to the ventral side, and the two lateral edges of the posteriormost primitive streak form the ventral ectodermal ridge (VER) (Goldman et al., 2000; Gruneberg, 1956). Consistent with the origin of this structure in primitive streak epiblast, YFP+ cells labelled at E7.5 were present in the VER at E10.5 (Figure 4Dc). These findings indicate that cells that expressed *Nkx1.2* at E7.5 or their descendants persist in the caudal end of the embryo at E10.5 and continue to generate largely neural tissue and paraxial mesoderm.

### Nkx1.2CreER^T2^ labels Sox2/Bra co-expressing cells

In the developing embryo, there is evidence for a neuromesodermal progenitor (NMp) cell population in the CLE and the NSB, where *Nkx1.2* is expressed, that contributes to neural and paraxial mesoderm lineages. This is based on retrospective clonal lineage analysis (Tzouanacou et al., 2009), grafting of cell groups in mouse (Cambray and Wilson, 2002, 2007; Wymeersch et al., 2016), and labelling of small cell groups in chick embryos (Brown and Storey, 2000). NMps made in vitro also express *Nkx1.2*. (Gouti et al., 2014). A key hallmark of NMps is the co-expression of the early neural and mesodermal markers Sox2 and Bra (Garriock et al., 2015; Tsakiridis et al., 2014; Wymeersch et al., 2016). To establish whether Nkx1.2CreER^T2^/YFP-labelled cells include NMps, we sought to identify YFP-labelled cells that co-express Sox2 and Bra by immunocytochemistry. Nkx1.2CreER^T2^ /YFP embryos exposed to tamoxifen at E7.5 and assessed at E8.5 (n=7 embryos) show that most CLE YFP+ cells in which *Nkx1.2* expression is driven express Sox2, and that a subset of such cells located in the CLE/NSB co-express Bra (Figure 5A). In embryos examined a day later, at E9.5 (n=4 embryos), YFP^+^ cells in the CLE/NSB continue to co-express Sox2 and Bra (Figure 5B). YFP^+^ cells, however, are absent from the caudal end of the notochord, where cells express high levels of Bra protein (Wymeersch et al., 2016)(Figure 5B). Similarly, at E10.5 (n=7 embryos), YFP+ cells labelled at E7.5 and/or their progeny co-express Sox2 and Bra in the neuroectoderm of the CNH region (Figure 5C), where grafting experiments indicate that NMps reside at this stage (Cambray and Wilson, 2002).

**Figure 5.**
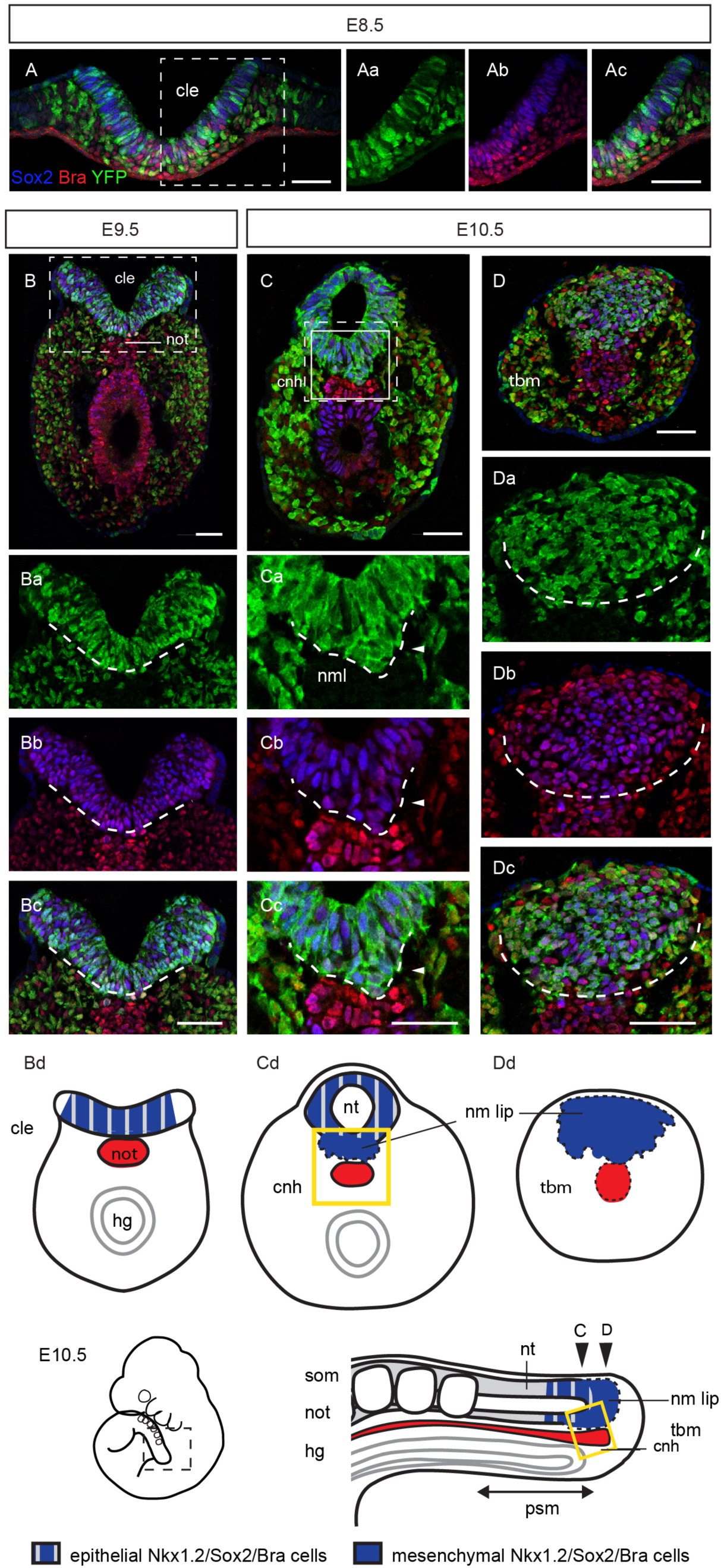
A subset of *Nkx1.2*-expressing cells or their progeny co-express Sox2 and Bra. Timed-pregnant Nkx1.2CreER^T2^/YFP mice received tamoxifen at E7.5 and embryos were analysed at (A) E8.5, (B) E9.5, and (C,D) E10.5. Transverse sections (TS) through the caudal end of these embryos were immunostained against Sox2 (blue), Bra (red), and YFP (green). (A) TS through the anterior primitive streak region of an E8.5 embryo. (Aa-c) Higher magnification of the CLE (cle) as indicated in A (dashed area). (B) TS through the CLE/node-streak border region of a E9.5 embryo. (Ba-c) Higher magnification of the CLE/node-streak border as indicated in B (dashed area). Sox2/Bra were co-expressed only in the CLE. (C) TS through the CNH region in an E10.5 embryo. (Ca-c) Higher magnification of the CNH as indicated in C (dashed area). YFP-labelled Sox2/Bra co-expressing cells extended ventrally from the neural tube above the end of the notochord at the CNH. Here, we propose the term neuromesodermal lip (nml) to define this region. (D) TS caudal to the CNH region at E10.5, through the tailbud mesenchyme (tbm). (Ca-c) Higher magnification of the tailbud mesenchyme as indicated in D. n = > 6 embryos per stage. Scale bars, 50 μm. (Bd, Cd, Dd) Schematics showing the location of Nkx1.2/Sox2//Bra-expressing cells in the developing caudal end of the mouse embryo at E9.5 (Bd) and E10.5 (Cd, Dd) in TS. Below, lateral view of the E10.5 tail bud. Nkx1.2/Sox2//Bra-expressing cells with epithelial morphology are coloured in blue/grey stripes. These cells are characteristic of the neuroectoderm/epiblast. Mesenchymal Nkx1.2/Sox2/Bra-expressing cells, in blue, are characteristic of the neuromesodermal lip, contiguous with caudal-most neural tube. Labels are as in Figure 1.

At E10.5, for the first time, Sox2/Bra co-expressing cells are also found in midline mesenchyme that now extends ventrally from the neural tube above the end of the notochord component of the CNH and also caudal to this into the tailbud mesenchyme (Figure 5C, D). Given this co-expression of neural and mesodermal genes, we propose here to name this medial mesenchymal cell population the neuromesodermal lip (NML) (Figure 5Cd, Dd). Importantly, this medial cell group coincides with the novel *Nkx1.2* domain identified above (Figure 1Ib-c), raising the possibility that these mesenchyme cells, which exhibit neural gene expression, are not only the source of neural tissue generated by secondary neurulation, but may also give rise to new paraxial mesoderm in the tailbud.

### Nkx1.2CreER^T2^ cells continue to make neural and mesodermal tissue in the developing tailbud

To test whether late E10.5 *Nkx1.2*-expressing cells indeed retain NM potential, timed-pregnant Nkx1.2CreER^T2^/YFP mice were exposed to tamoxifen at E10.5 and embryos assessed a day later, at E11.5. In these embryos, labelled cells contributed to neural and mesodermal tissues, but were now not found in the hindgut or surface ectoderm, (n=13 embryos, 9 assessed in sections) (Figure 6). Labelled cells in the spinal cord were found from immediately below the hindlimb (opposite somite ~36) to the tailbud, while labelled cells in the paraxial mesoderm were found only in the newly emerged presomitic mesoderm (compare Figure 6a and 6b–d). This different distribution of labelled cells along the rostrocaudal axis probably reflects the broader expression of *Nkx1.2* in the caudal neural tube (Figure 1I, K). The presence of labelled cells within the E11.5 tailbud mesenchyme further raises the possibility that some cells are retained here and may therefore progressively give rise to neural and mesodermal tissues in the developing tailbud. As some of these cells co-express Sox2/Bra and labelled progeny are found in paraxial mesoderm as well as neural tissue (Figure 6), it is tempting to speculate that these *Nkx1.2*-expressing cells constitute a reservoir of bipotent NMps for secondary body formation. However, without single cell labelling we cannot rule out the possibility that individual cells (or even spatially distinct cell groups) within the *Nkx1.2* domain generate neural or mesodermal lineages.

**Figure 6.**
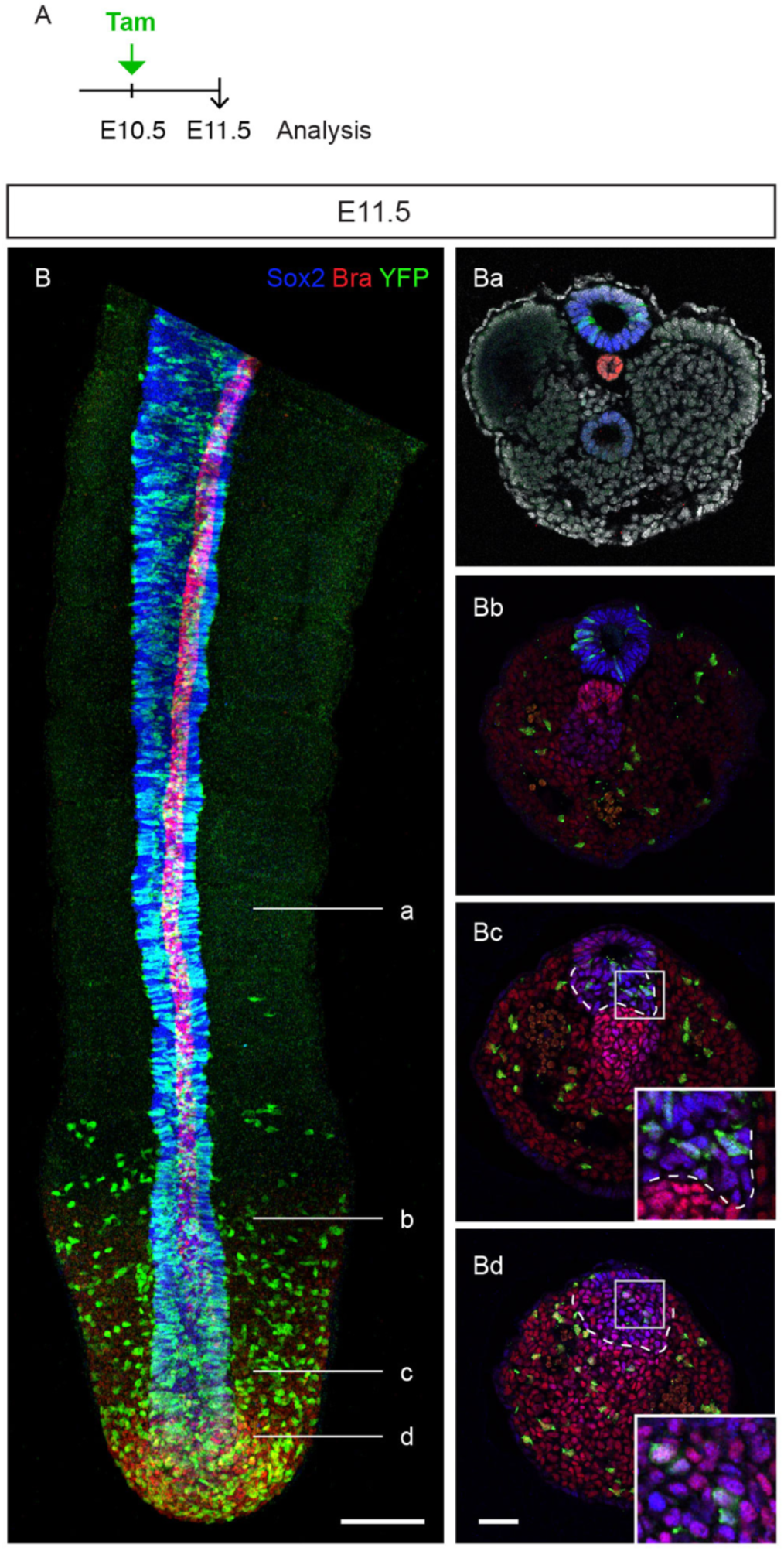
E.10.5 *Nkx1.2*-expressing cells continue to generate neural and mesodermal tissues. (A) Timed-pregnant Nkx1.2CreER^T2^/YFP mice received tamoxifen at E10.5 and embryos were analysed at E11.5. (B) Confocal MIP of the caudal end of the tail of an E11.5 embryo labelled with antibodies against YFP (green), Sox2 (blue), and Bra (red). Dorsal view. (Aa-d) Transverse sections through the levels indicated in B. YFP^+^ cells formed secondary neural tube and presomitic mesoderm. YFP+ cells were also found in the neuromesodermal lip (Bc) and contiguous caudal mesenchyme (Bd). Scale bars, 50 μm.

Overall, these data document the restriction of caudal lateral epiblast cell fates, but support the possibility that bipotent neuromesodermal progenitors persist to the end of axis elongation and contribute to neural tissue formed by secondary neurulation.

## Materials and methods

### Mice and tamoxifen administration

To make a tamoxifen stock solution, tamoxifen powder (Sigma T5648) was dissolved in vegetable oil to a final concentration of 40 mg/ml and sonicated to bring to solution. The tamoxifen stock solution was stored at ‐20 °C for up to 3 months. The morning of the plug was considered (E0.5). At various stages of pregnancy, Nkx1.2CreER^T2^ /YFP females were given a single 200μl dose of tamoxifen (of the 40 mg/ml stock) via oral gavage. Mice were monitored for 6 hours and when required sacrificed following schedule 1 of the Animals (Scientific Procedures) Act of 1986. Embryos were dissected in cold PBS and fixed in cold 4% PFA for 2 hours. Gastrula embryos were staged according to (Downs and Davies, 1993) and at later stages by standard morphological criteria.

### Genotyping

Genotyping by standard methods was performed to maintain the homozygous line using the following PCR conditions: 95°C, 5 min and then 95°C, 30s; 60°C, 30 s; 72°C, 1 min for 35 cycles followed by 72°C, 10 min.

A DNA quality control and a test reaction were carried out in parallel for the KI allele, the wild-type (WT) allele, and the Flpe deleter (TG) using the following primer pairs: KI primer 1: 5’ACGTCCAGACACAGCATAGG 3’, primer 2: 5’TCACTGAGCAGGTGTTCAGG 3’ (fragment size 279 bp); QC primer 3: 5’GAGACTCTGGCTACTCATCC 3’; primer 4: 5’CCTTCAGCAAGAGCTGGGGAC 3’ (fragment size 585 bp); WT primer 5: 5’CAAGGTTTATTGGTAGCCTGG 3’, primer 6: 5’TGAGCCAGTCAGAGTTGTGG 3’ (fragment size 176 bp); QC primer 7: 5’GTGGCACGGAACTTCTAGTC, primer 8: 5’CTTGTCAAGTAGCAGGAAGA 3’ (fragment size 335 bp); TG primer 9: 5’GGCAGAAGCACGCTTATCG 3’, primer 10: 5’GACAAGCGTTAGTAGGCACAT 3’ (fragment size 343 bp); QC primer 3 as above, QC P4 as above (fragment size 585 bp).

### Antibody staining and imaging

Embryos were permeablised by dehydration in methanol (25% methanol/PBS, 50% methanol/PBS, 100% methanol), bleached in 3% H_2_O2/methanol, rehydrated in PBS, and incubated with anti-GFP antibodies to enhance endogenous YFP signal either as whole embryos or on cross sections. Whole embryos were pre-blocked in PBS/0.1% Tween-20 (PBST) and 10% normal donkey serum (NDS) for 4 hours, incubated overnight with primary antibodies in PBST/NDS (1:500), washed extensively in PBST, and incubated with secondary antibodies (1:500) overnight. Embryos for cryosectioning were equilibrated in 30% sucrose/PBS, mounted in agar blocks (1.5% agar/5% sucrose/PBS), and frozen on dry ice. Sections of 16 μm were cut on a Leica CM1900 cryostat and mounted on poly-lysine coated slides. Whole embryos were imaged on a Leica DM fluorescence dissecting microscope and a subset of early embryos and isolated whole tails on a Leica SP8 confocal microscope. Sections were scored on a Leica DB fluorescence microscope or with a DeltaVision Imaging system, and images acquired on a Leica SP8 confocal microscope. Antibodies used were: chicken anti-GFP (Abcam, ab13970), goat anti-GFP (Abcam, ab6673), rabbit anti-SOX2 (Millipore, AB5603), goat anti-SOX2 (Immune Systems, GT15098), goat anti-T (R&D, AF2085), donkey anti-chicken Alexa Fluor 488 (Abcam, ab150173), donkey anti-goat Alexa Fluor 488 (Life Technologies, A11055), donkey anti-rabbit Alexa Fluor 568 (Life Technologies, A10042), donkey anti-goat Alexa Fluor 647 (Life Technologies, A21477).

### RNA in situ hybridization

Standard methods were used to carry out mRNA in situ hybridisation in wild type CD1 embryos (Wilkinson and Nieto, 1993). *Nkx1.2* plasmid was kindly provided by Frank Schubert. This probe includes the homeobox domain and the 3’ half of the gene (nucleotides 504-1057).

### Ethics statement

All procedures using animals were performed in accordance with UK and EU legislation and guidance on animal use in bioscience research. The work was carried out under the UK project license 60/4454 and was subjected to local ethical review.

## Acknowledgements

We thank the University of Dundee WBRU-TG for technical assistance and advice, Frank Schubert for kindly providing the *Nkx1.2* plasmid probe, and Val Wilson and Moises Mallo for prompt and insightful comments on the manuscript. Creation of and initial work with the Nkx1.2CreER^T2^ line was supported MRC grant G1100552 (P.A.H. and K.G.S.) and also by a Wellcome Trust Senior Investigator Award to KGS (A.R.A) (WT102817).

## Author contributions

Experiments were carried out and analysed by PAH, ARA and KGS. ARA and KGS wrote the paper.

The authors declare no competing interests.

## References

Bae, Y.K., Shimizu, T., Muraoka, O., Yabe, T., Hirata, T., Nojima, H., Hirano, T., and Hibi, M. (2004). Expression of sax1/nkx1.2 and sax2/nkx1.1 in zebrafish. Gene Expr Patterns 4, 481–486.

Bate, M. (1990). The embryonic development of larval muscles in Drosophila. Development (Cambridge, England) 110, 791–804.

Bober, E., Baum, C., Braun, T., and Arnold, H.H. (1994). A novel NK-related mouse homeobox gene: expression in central and peripheral nervous structures during embryonic development. Dev Biol 162, 288–303.

Brown, J.M., and Storey, K.G. (2000). A region of the vertebrate neural plate in which neighbouring cells can adopt neural or epidermal cell fates. Current Biology 10, 869–872.

Cambray, N., and Wilson, V. (2002). Axial progenitors with extensive potency are localised to the mouse chordoneural hinge. Development (Cambridge, England) 129, 4855–4866.

Cambray, N., and Wilson, V. (2007). Two distinct sources for a population of maturing axial progenitors. Development (Cambridge, England) 134, 2829–2840.

Delfino-Machin, M., Lunn, J.S., Breitkreuz, D.N., Akai, J., and Storey, K.G. (2005). Specification and maintenance of the spinal cord stem zone. Development (Cambridge, England) 132, 4273–4283.

Diez del Corral, R., Breitkreuz, D.N., and Storey, K.G. (2002). Onset of neuronal differentiation is regulated by paraxial mesoderm and requires attenuation of FGF signalling. Development (Cambridge, England) 129, 1681–1691.

Diez del Corral, R., Olivera-Martinez, I., Goriely, A., Gale, E., Maden, M., and Storey, K. (2003). Opposing FGF and retinoid pathways control ventral neural pattern, neuronal differentiation, and segmentation during body axis extension. Neuron 40, 65–79.

Dohrmann, C., Azpiazu, N., and Frasch, M. (1990). A new Drosophila homeo box gene is expressed in mesodermal precursor cells of distinct muscles during embryogenesis. Genes Dev 4, 2098–2111.

Downs, K.M., and Davies, T. (1993). Staging of gastrulating mouse embryos by morphological landmarks in the dissecting microscope. Development (Cambridge, England) 118, 1255–1266.

Forlani, S., Lawson, K.A., and Deschamps, J. (2003). Acquisition of Hox codes during gastrulation and axial elongation in the mouse embryo. Development (Cambridge, England) 130, 3807–3819.

Garriock, R.J., Chalamalasetty, R.B., Kennedy, M.W., Canizales, L.C., Lewandoski, M., and Yamaguchi, T.P. (2015). Lineage tracing of neuromesodermal progenitors reveals novel Wnt-dependent roles in trunk progenitor cell maintenance and differentiation. Development (Cambridge, England) 142, 1628–1638.

Goldman, D.C., Martin, G.R., and Tam, P.P. (2000). Fate and function of the ventral ectodermal ridge during mouse tail development. Development (Cambridge, England) 127, 2113–2123.

Gouti, M., Tsakiridis, A., Wymeersch, F.J., Huang, Y., Kleinjung, J., Wilson, V., and Briscoe, J. (2014). In vitro generation of neuromesodermal progenitors reveals distinct roles for wnt signalling in the specification of spinal cord and paraxial mesoderm identity. PLoS Biol 12, e1001937.

Gruneberg, H. (1956). A ventral ectodermal ridge of the tail in mouse embryos. Nature 177, 787–788.

Hamburger, H., and Hamilton, H.L. (1951). A series of normal stages in the development of the chick embryo. J Exp Morphol 88, 49–92.

Henrique, D., Abranches, E., Verrier, L., and Storey, K.G. (2015). Neuromesodermal progenitors and the making of the spinal cord. Development (Cambridge, England) 142, 2864–2875.

Kumar, S., and Duester, G. (2014). Retinoic acid controls body axis extension by directly repressing Fgf8 transcription. Development (Cambridge, England) 141, 2972–2977.

Lawson, K.A., Meneses, J.J., and Pedersen, R.A. (1991). Clonal analysis of epiblast fate during germ layer formation in the mouse embryo. Development (Cambridge, England) 113, 891–911.

Lawson, K.A., and Pedersen, R.A. (1992). Clonal analysis of cell fate during gastrulation and early neurulation in the mouse. Ciba Found Symp 165, 3-21; discussion 21-26.

Marikawa, Y., Tamashiro, D.A., Fujita, T.C., and Alarcon, V.B. (2009). Aggregated P19 mouse embryonal carcinoma cells as a simple in vitro model to study the molecular regulations of mesoderm formation and axial elongation morphogenesis. Genesis 47, 93–106.

Olivera-Martinez, I., Harada, H., Halley, P.A., and Storey, K.G. (2012). Loss of FGF-dependent mesoderm identity and rise of endogenous retinoid signalling determine cessation of body axis elongation. PLoS Biol 10, e1001415.

Rangini, Z., Frumkin, A., Shani, G., Guttmann, M., Eyal-Giladi, H., Gruenbaum, Y., and Fainsod, A. (1989). The chicken homeo box genes CHox1 and CHox3: cloning, sequencing and expression during embryogenesis. Gene 76, 61–74.

Sasai, N., Kutejova, E., and Briscoe, J. (2014). Integration of Signals along Orthogonal Axes of the Vertebrate Neural Tube Controls Progenitor Competence and Increases Cell Diversity. PLoS Biol 12, e1001907.

Schoenwolf, G.C. (1984). Histological and ultrastructural studies of secondary neurulation in mouse embryos. Am J Anat 169, 361–376.

Schubert, F.R., Fainsod, A., Gruenbaum, Y., and Gruss, P. (1995). Expression of a novel murine homeobox gene Sax-1 in the developing nervous system. Mechanisms of Development 51, 99–114.

Schubert, F.R., and Lumsden, A. (2005). Transcriptional control of early tract formation in the embryonic chick midbrain. Development (Cambridge, England) 132, 1785–1793.

Simon, R., and Lufkin, T. (2003). Postnatal lethality in mice lacking the Sax2 homeobox gene homologous to Drosophila S59/slouch: evidence for positive and negative autoregulation. Mol Cell Biol 23, 9046–9060.

Sirbu, I.O., and Duester, G. (2006). Retinoic-acid signalling in node ectoderm and posterior neural plate directs left-right patterning of somitic mesoderm. Nat Cell Biol 8, 271–277.

Snow, M.H.L. (1977). Gastrulation in the mouse: Growth and regionalization of the epiblast. Development (Cambridge, England) 42, 293–303.

Spann, P., Ginsburg, M., Rangini, Z., Fainsod, A., Eyal Giladi, H., and Gruenbaum, Y. (1994). The spatial and temporal dynamics of Sax1 (CHox3) homeobox gene expression in the chick’s spinal cord. Development (Cambridge, England) 120, 1817–1828.

Srinivas, S., Watanabe, T., Lin, C.S., William, C.M., Tanabe, Y., Jessell, T.M., and Costantini, F. (2001). Cre reporter strains produced by targeted insertion of EYFP and ECFP into the ROSA26 locus. BMC developmental biology 1, 4.

Tam, P.P., and Beddington, R.S. (1987). The formation of mesodermal tissues in the mouse embryo during gastrulation and early organogenesis. Development (Cambridge, England) 99, 109–126.

Tamashiro, D.A., Alarcon, V.B., and Marikawa, Y. (2012). Nkx1-2 is a transcriptional repressor and is essential for the activation of Brachyury in P19 mouse embryonal carcinoma cell. Differentiation 83, 282–292.

Tanaka, S.S., Kojima, Y., Yamaguchi, Y.L., Nishinakamura, R., and Tam, P.P. (2011). Impact of WNT signaling on tissue lineage differentiation in the early mouse embryo. Dev Growth Differ 53, 843–856.

Tsakiridis, A., Huang, Y., Blin, G., Skylaki, S., Wymeersch, F., Osorno, R., Economou, C., Karagianni, E., Zhao, S., Lowell, S., et al. (2014). Distinct Wnt-driven primitive streak-like populations reflect in vivo lineage precursors. Development (Cambridge, England) 141, 1209–1221.

Tsakiridis, A., and Wilson, V. (2015). Assessing the bipotency of in vitro-derived neuromesodermal progenitors. F1000Research 4, 100.

Turner, D.A., Rue, P., Mackenzie, J.P., Davies, E., and Martinez Arias, A. (2014). Brachyury cooperates with Wnt/beta-catenin signalling to elicit primitive-streak-like behaviour in differentiating mouse embryonic stem cells. BMC Biol 12, 63.

Tzouanacou, E., Wegener, A., Wymeersch, F.J., Wilson, V., and Nicolas, J.F. (2009). Redefining the progression of lineage segregations during mammalian embryogenesis by clonal analysis. Dev Cell 17, 365–376.

Wilkinson, D.G., and Nieto, M.A. (1993). Detection of messenger RNA by in situ hybridization to tissue sections and whole mounts. Methods Enzymol 225, 361–373.

Wilson, V., and Beddington, R.S. (1996). Cell fate and morphogenetic movement in the late mouse primitive streak. Mech Dev 55, 79–89.

Wilson, V., Olivera-Martinez, I., and Storey, K.G. (2009). Stem cells, signals and vertebrate body axis extension. Development (Cambridge, England) 136, 1591–1604.

Wymeersch, F.J., Huang, Y., Blin, G., Cambray, N., Wilkie, R., Wong, F.C., and Wilson, V. (2016). Position-dependent plasticity of distinct progenitor types in the primitive streak. Elife 5.

